# Exon probe sets and bioinformatics pipelines for all levels of fish phylogenomics

**DOI:** 10.1101/2020.02.18.949735

**Authors:** Lily C. Hughes, Guillermo Ortí, Hadeel Saad, Chenhong Li, William T. White, Carole C. Baldwin, Keith A. Crandall, Dahiana Arcila, Ricardo Betancur-R.

## Abstract

Exon markers have a long history of use in phylogenetics of ray-finned fishes, the most diverse clade of vertebrates with more than 35,000 species. As the number of published genomes increases, it has become easier to test exons and other genetic markers for signals of ancient duplication events and filter out paralogs that can mislead phylogenetic analysis. We present seven new probe sets for current target-capture phylogenomic protocols that capture 1,104 exons explicitly filtered for paralogs using gene trees. These seven probe sets span the diversity of teleost fishes, including four sets that target five hyper-diverse percomorph clades which together comprise *ca*. 17,000 species (Carangaria, Ovalentaria, Eupercaria, and Syngnatharia + Pelagiaria combined). We additionally included probes to capture exon markers that have been commonly used in fish phylogenetics, despite some being flagged for paralogs, to facilitate integration of old and new molecular phylogenetic matrices. We tested these probes and merged new exon-capture sequence data into an existing data matrix of 1,105 exons and 300 ray-finned fish species. We provide an optimized bioinformatics pipeline to assemble exon capture data from raw reads to alignments for downstream analysis. We show that loci with known paralogs are at risk of assembling duplicated sequences with target-capture, but we also assembled many useful orthologous sequences. These probe sets are a valuable resource for advancing fish phylogenomics because they can be easily extracted from increasingly available whole genome and transcriptome datasets, and also may be integrated with existing PCR-based exon and mitochondrial datasets.

## Introduction

Exon markers have played a pivotal role in resolving phylogenetic relationships among ray-finned fishes (Li *et al*. 2007; Near *et al*. 2012; Betancur-R. *et al*. 2013; Betancur-R *et al*. 2017; Hughes *et al*. 2018; Rabosky *et al*. 2018). Identification of these exons has typically involved the comparison of a small number of fish model genomes. For example, a suite of 154 single-copy exons conserved enough for PCR amplification was identified by Li *et al*. (2007). This study used a reciprocal BLAST approach on two genomes, the pufferfish *Takifugu rubripes* and the zebrafish *Danio rerio*, with 10 exons initially optimized with PCR primers for sequencing (Li *et al*. 2007). These exons demonstrated their utility for resolving previously enigmatic relationships among fishes (Li *et al*. 2008), and were the basis for largescale reappraisals of the ray-finned fish Tree of Life (Near *et al*. 2012; Betancur-R. *et al*. 2013), phylogenetic analysis of the large clade of the “spiny-ray” acanthomorph fishes (Near *et al*. 2013b), and new phylogenetic classifications based on sequence data for more than 2,000 fish species (Betancur-R. *et al*. 2013; Betancur-R *et al*. 2017). Most recently, these exons formed a large part of a supermatrix analysis that utilized sequence data for more than 11,000 ray-finned fish species available through GenBank in one of the largest analyses to date (Rabosky *et al*. 2018).

The advent of high-throughput sequencing technologies has drastically increased the number of loci systematists can harness for their groups of interest. Sequence capture based on single-stranded RNA probes that enrich genomic DNA libraries for conserved molecular markers have revolutionized phylogenomics (Faircloth *et al*. 2012; Lemmon *et al*. 2012), allowing cost-effective sequencing of hundreds or thousands of markers for many taxa. Ultraconserved Elements (UCEs) (Faircloth *et al*. 2012) have been used for fish phylogenomics (Faircloth *et al*. 2013, 2020; Harrington *et al*. 2016; Longo *et al*. 2017; Chakrabarty *et al*. 2017; Alfaro *et al*. 2018; Roxo *et al*. 2019; Friedman *et al*. 2019), as have Anchored Hybrid Enrichment (AHE) approaches (Lemmon *et al*. 2012; Eytan *et al*. 2015; Stout *et al*. 2016; Dornburg *et al*. 2017; Irisarri *et al*. 2018). But exon markers have desirable properties for phylogenomics that other approaches may lack. They are relatively easy to align, and a number of software programs have been developed for reading frame-aware alignment (Abascal *et al*. 2010; Ranwez *et al*. 2011, 2018), avoiding potential homology errors with UCE flanking regions that are more difficult to align (Edwards *et al*. 2017). Both protein and nucleotide sequences can be used for phylogenetic inference, making them useful for deep (Hughes *et al*. 2018) and shallow phylogenetic scales (Rincon-Sandoval *et al*. 2019). Exon markers are also easy to integrate with both genomic and transcriptomic data, with sequences being produced and archived for a variety of studies from comparative genomics to gene expression analysis that can provide resources for systematists to increase taxon sampling without incurring additional costs.

Since exon markers tend to be more variable across the target region than UCEs or AHE, two rounds of *in vitro* hybridization are optimal for their sequence capture protocols (Li *et al*. 2013). This improvement in laboratory techniques has resulted in a number of studies using sequence capture to target exons for fish phylogenomics (Ilves & López-Fernández 2014; Li *et al*. 2015; Song *et al*. 2017; Arcila *et al*. 2017; Ilves *et al*. 2017; Kuang *et al*. 2018; Straube *et al*. 2018; Betancur-R. *et al*. 2019; Rincon-Sandoval *et al*. 2019; Yin *et al*. 2019). The increase in genomic resources for fishes has allowed for the comparison of a larger number of genomes for probe design (Li *et al*. 2012), and ultimately eight ray-finned fish genomes were used to identify >17,000 ‘single-copy’ exons (Song *et al*. 2017) using a modification of the reciprocal BLAST approach of Li *et al*. (2007). A subset (4,434) of these were optimized for capture across all ray-finned fishes (Jiang *et al*. 2019). Other sets of exon probes have been designed to target more specific groups including cichlids (Ilves & López-Fernández 2014), and otophysans (Arcila *et al*. 2017).

The evolutionary history of vertebrate genomes is complicated by two ancient Whole-Genome Duplication (WGD) events, and an additional WGD event in the ancestor to all modern teleosts (Vandepoele *et al*. 2004; Dehal & Boore 2005). While many duplicated gene copies were lost shortly after the teleost WGD (Inoue *et al*. 2015), up to a quarter of genes in teleost genomes have paralogs as a consequence of this event (Braasch *et al*. 2015). Phylogenetic inference relies on the analysis of orthologs, but loci implemented so far for analyses were based on the comparison of a limited number of model genomes and some threshold of similarity to define them as “single-copy” (Li *et al*. 2007). A recent study implementing an explicit tree-based filtering method to test for orthology revealed that one third of the ‘single-copy’ exons >200 bp in length identified by Jiang *et al*. (2019) were affected by paralogy, potentially biasing tree inference (Hughes *et al*. 2018). A set of 1,105 exons free of vertebrate and teleost WGD-derived paralogs identified in this study resolved the phylogeny with confidence for more than 300 species of ray-finned fishes. Other markers used for phylogenomic studies such as UCEs (Faircloth *et al*. 2013), AHE loci (Lemmon *et al*. 2012) and exons (Ilves & López-Fernández 2014; Song *et al*. 2017; Arcila *et al*. 2017; Jiang *et al*. 2019) also have not been explicitly tested for paralogy using approaches based on gene trees.

Here we present a new experimental protocol to obtain sequence data for a set of exons filtered for paralogs (Hughes *et al*. 2018) across the diversity of fishes. We provide probe sets for target capture that are designed to enrich genomic libraries for different taxonomic groups, from the early branching teleosts to the major groups within percomorphs, the massive radiation comprising more than 17,000 species. These probe sets target the same loci, but the specific sequences of the probes are tailored to capture more efficiently within taxonomic brackets. We have also included probes for other legacy exon markers (e.g., Lopez *et al*. 2004; Lovejoy *et al*. 2004; Dettai & Lecointre 2005; Li *et al*. 2007) and mitochondrial DNA (mtDNA) loci that have been sequenced for a large number of fishes through PCR-Sanger sequencing methods to facilitate integration of new high throughput sequencing results with existing phylogenetic datasets. We designed a new bioinformatics pipeline that wraps aTRAM 2.0 (Allen *et al*. 2017) with other software packages and python scripts to assemble and filter sequence alignments from Illumina reads.

## Methods

### Nuclear Exon Probes

Sequences for probe design came from exon alignments derived from a database of 303 bony fish genomes and transcriptomes (Sun *et al*. 2016; Hughes *et al*. 2018). Briefly, the EvolMarkers pipeline (Li *et al*. 2012, 2015; Jiang *et al*. 2019) was used to identify 1,721 single-copy exons in eight ray-finned fish genomes (*Lepisosteus oculatus*, *Anguilla japonica, Danio rerio, Gadus morhua, Oreochromis niloticus, Oryzias latipes, Tetraodon nigroviridis*, and *Gasterosteus aculeatus*). These exons were mined from 295 other genomes and transcriptomes using nhmmer (Wheeler & Eddy 2013) in HMMER v3.1b2, and exons with paralogs were filtered by testing for duplications in gene trees via topology tests (see Hughes *et al*. 2018 for full details).

A total of 1,105 exons were retained after filtering for loci with paralogs. We generated seven probe sets for these exons based on different underlying references for our target groups, following the classification of Betancur-R. *et al*. (2017). These seven target teleost groups included (i) Elopomorpha (~1000 species, including true eels and tarpons) (Figure 1); (ii) early branching teleosts from Osteoglossomorpha (bonytongues) to Myctophiformes (lanternfishes)— hereafter paraphyletic “Backbone 1” (Figure 1); (iii) Acanthomorphata (from paracanthopterygians (e.g. cods, oarfish) to Anabantaria (e.g. swamp eels, gouramies)— hereafter paraphyletic “Backbone 2”) (Figure 2); and four specific sets aimed for some of the most species-rich clades of Percomorphaceae, including (iv) Carangaria (~1,100 species, including flatfishes and jacks) (Figure 3); (v) Ovalentaria (~5,600 species, including clownfishes, cichlids, flying fishes) (Figure 2); (vi) Eupercaria (~6,800 species, including surgeonfishes, pufferfishes, and groupers) (Figure 3); and (vii) Syngnatharia-Pelagiaria (~1,000 species, including tunas, seahorses, and pipefishes) (Figure 3). The large freshwater Otophysa clade (>10,000 species, including catfishes, knifefishes, and tetras) is not included in Backbone 1 (Figure 1), largely because it was targeted earlier by a more specific probe set designed for the clade by other exon-capture fish studies (Arcila *et al*. 2017; Betancur-R. *et al*. 2019), though 143 exons are shared between the two. We designed probe sets for different subsets of taxa from these 1,105 alignments that initially consisted of 303 species that span the diversity of bony fishes (Hughes *et al*. 2018), as explained above. Each of the seven probe set references were comprised four to eight of the most phylogenetically-distant taxa in the target clade, depending on the phylogenetic breadth the probe set covers (Table 1). We ranked preferred taxa within each of these groups to form the basis for the probe set references (Table 1), and if all preferred taxa were missing from a group, we took the next longest sequence in the alignment for the clade of interest. This means that some exons may have unique taxa representing them in their reference set. One particularly long exon included highly divergent sequences that were difficult to align and was ultimately excluded from the final probe sets (a total of 1,104 target exons remained).

**Figure 1:**
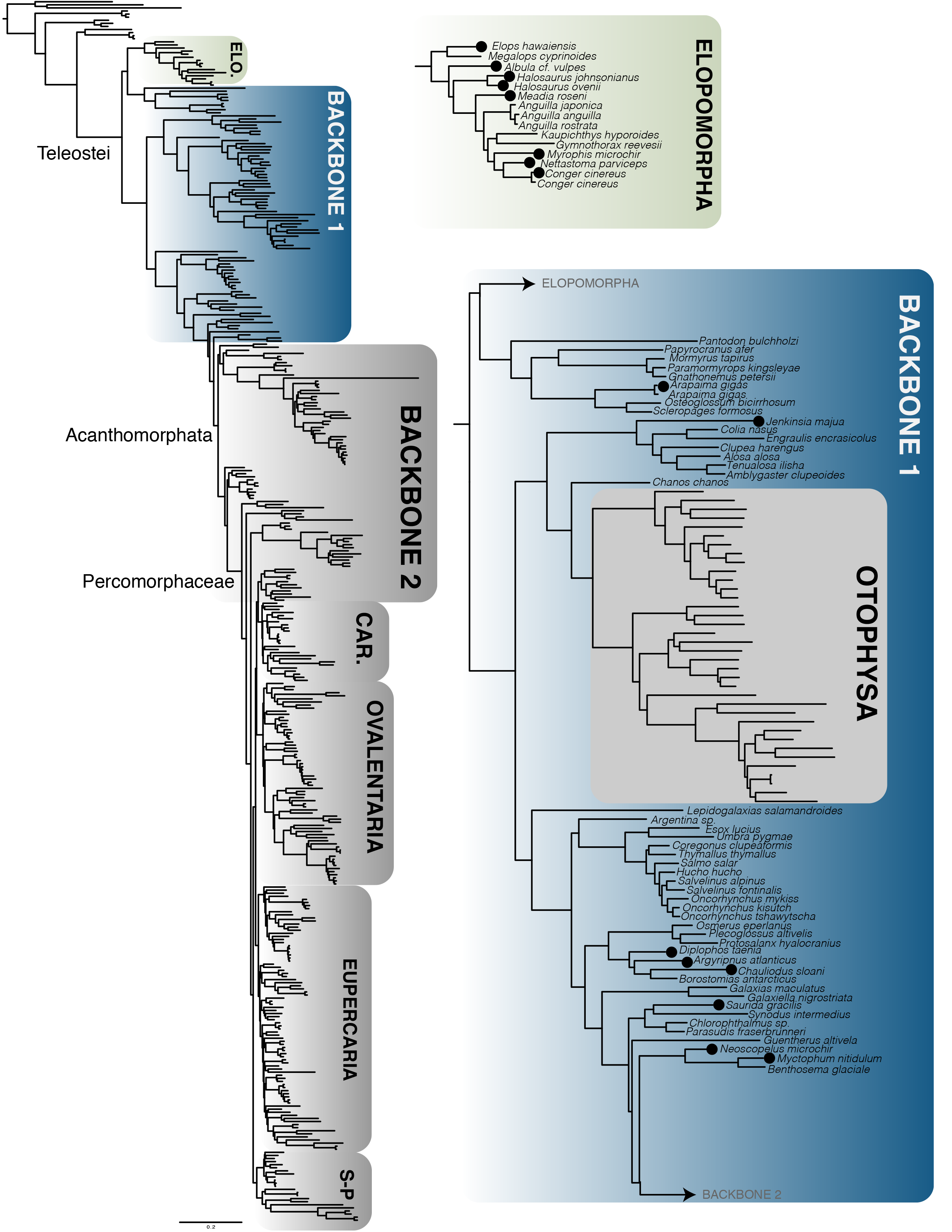
Position of Elopomorpha and Backbone 1 probe sets in the Maximum likelihood analysis of protein from 394 fish taxa, combining genomes, transcriptomes, and exon capture. Newly sequenced taxa are represented with black dots at the tips. This clade has a specific probe set with a different but overlapping set of exons designed in an earlier study, Arcila *et al*. (2017), but is not specifically targeted by Backbone 1.

**Figure 2:**
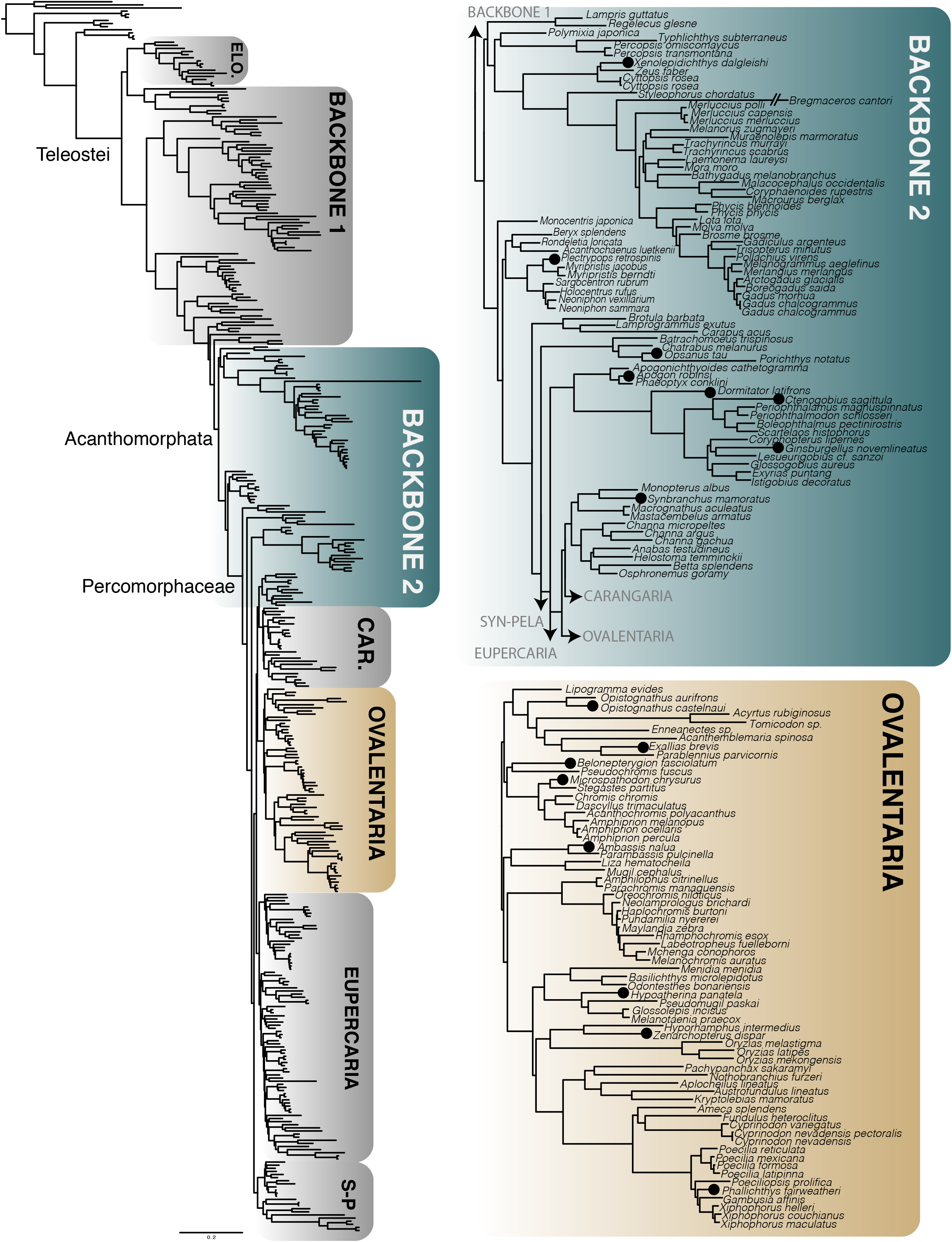
Position of Backbone 2 and Ovalentaria probe sets in the Maximum likelihood analysis of 394 fish taxa, combining genomes, transcriptomes, and exon capture. Newly sequenced taxa are represented with black dots at the tips.

**Figure 3:**
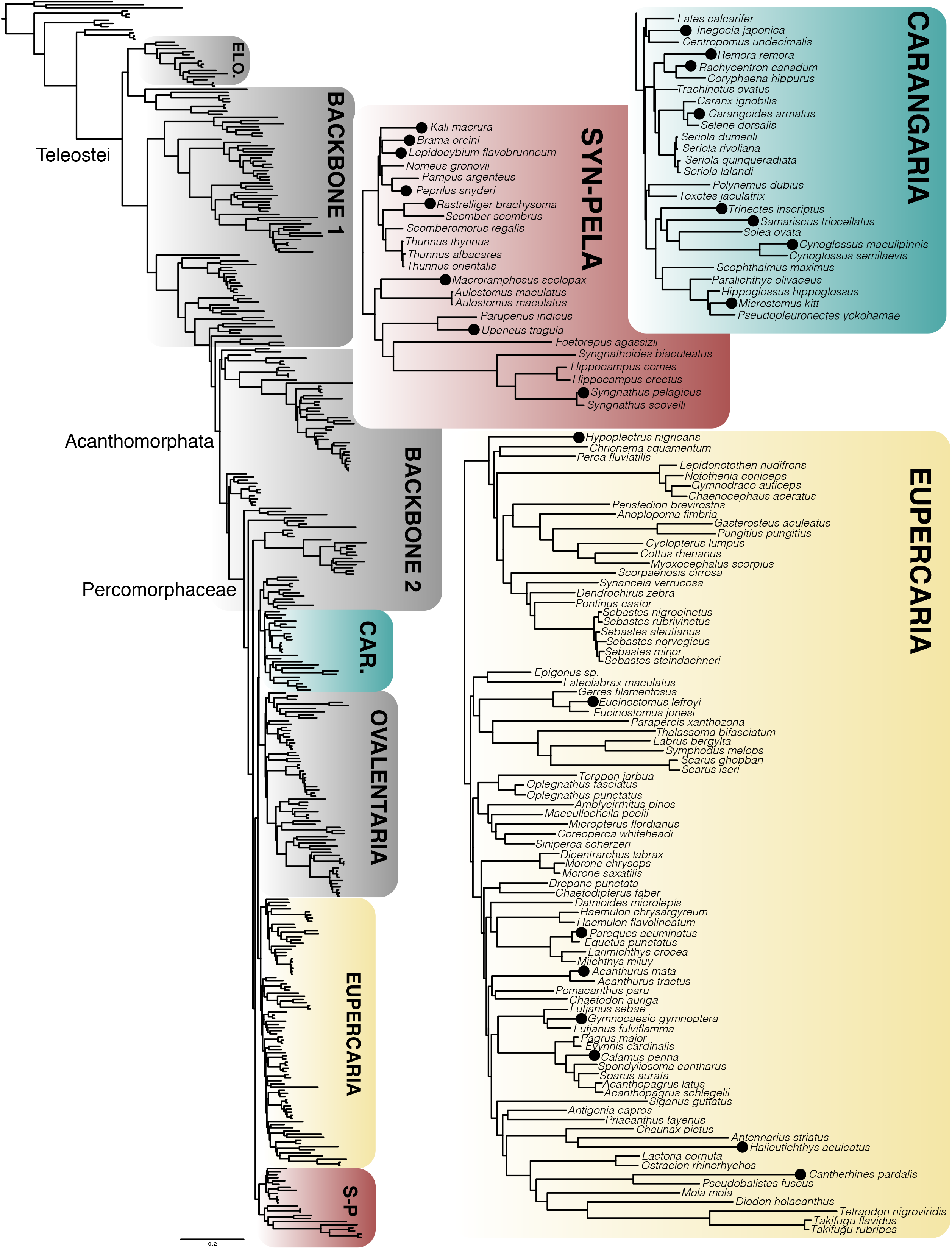
Position of Carangaria, Syngnatharia-Pelagiaria (Syn-Pela) and Eupercaria probe sets in the Maximum Likelihood analysis of 394 fish taxa, combining genomes, transcriptomes, and exon capture. Newly sequenced taxa are represented with black dots.

**Table 1.** Seven probe sets designed for 1,104 conserved exon markers across teleost fishes.

| Probe Set Name | Lineages Included (Preferred lineage in bold) |
| --- | --- |
| Elopomorpha | 1. **Megalopidae**<br>2. **Muraenidae**<br>3. **Congridae**, Chlopsidae<br>4. **Anguillidae** |
| Backbone 1 | 1. **Osteoglossidae**, Pantodontidae<br>2. **Notopteridae**, Mormyridae |
|  | 3. **Engraulidae**, Clupeidae<br>4. **Galaxiidae**<br>5. **Argentinidae**<br>6. **Stomiidae**, Osmeridae, Plecoglossidae, Salangidae<br>7. **Synodontidae**, Chlorophthalmidae<br>8. **Myctophidae** |
| Backbone 2 | 1. **Zeidae**, Parazenidae<br>2. **Berycidae**, Stephanoberycidae, Rondeletiidae<br>3. **Holocentridae**<br>4. **Ophidiidae**<br>5. **Apogonidae**<br>6. **Gobiidae**<br>7. **Synbranchiiformes**, Anabantiformes |
| Syngnatharia-Pelagiaria | 1. **Syngnathidae**, Callionymidae<br>2. **Mullidae**, Aulostomidae<br>3. **Scombridae**<br>4. **Nomeidae**, Stromateidae |
| Carangaria | 1. **Coryphaenidae**, Carangidae<br>2. **Cynoglossidae**, Paralichthyidae, Pleuronectidae, Scophthalmidae, Soleidae<br>3. **Centropomidae**<br>4. **Polynemidae**, Toxotidae |
| Ovalentaria | 1. **Pseudomugilidae**, Melanotaeniidae, Atherinopsidae<br>2. **Aplocheilidae**, Nothobranchiidae, Rivulidae, Cyprinodontidae, Fundulidae, Poeciliidae<br>3. **Tripterygiidae**, Blenniidae, Chaenopsidae<br>4. **Gobiesocidae**<br>5. **Pomacentridae** |
| Eupercaria | 1. **Anoplopomatidae**, Channichthyidae, Cottidae, Gasterosteidae, Nototheniidae, Bathydraconidae, Percidae, Sebastidae<br>2. **Gerreidae**, Labridae, Scaridae, Pinguipedidae, Lateolabracidae, Epigonidae<br>3. **Tetraodontidae**, Molidae, Chaunacidae, Caproidae, Diodontidae, Antennariidae, Balistidae, Acanthuridae<br>4. **Lutjanidae**, Haemulidae, Chaetodontidae, Moronidae, Datnioididae, Ephippidae, Sciaenidae |

We included baits for nuclear markers popular in fish phylogenetics (hereafter referred to as ‘legacy’ markers) to better connect sequence datasets produced by targeted amplicon sequencing approaches (Bybee *et al*. 2011). Several of these widely used markers were already included as part of the “paralogy-tested” 1,105 exons from Hughes *et al*. (2018), including RAG1 (Lopez *et al*. 2004), RAG2 (Lovejoy *et al*. 2004), FICD (Li *et al*. 2011), PANX2, GCS1, GLYT (Li *et al*. 2007), VCPIP (Betancur-R. *et al*. 2013), and MLL (Dettai & Lecointre 2005). A total of 19 additional legacy markers that did not meet the paralogy filtering requirements were nonetheless included in the probe sets, mainly markers developed by the Euteleost Tree of Life Project: TBR1, MYH6, KIAA1239, PLAGL2, BTCHD1, RIPK4, SH3PX3, SIDKEY, SREB2, ZIC1, SVEP1, GPR61, SLC10A3, UBE3A, and UBE3A-like (Li *et al*. 2007, 2011; Betancur-R. *et al*. 2013). Additionally, baits were designed for the markers Rhodopsin (Chen *et al*. 2003), IRBP (Dettaï & Lecointre 2008), and RNF213 (Li *et al*. 2009a), which have been widely used in fish systematics. Due to the long sequences of MYH6 and KIAA1239 (>3,000 bp), references for these markers were shortened to the region typically amplified by PCR primers. The reference sequences used in bait design are available on GitHub (https://github.com/lilychughes/FishLifeExonCapture/tree/master/ProbeSets).

Probe sequences of 120 bp in length were initially generated with the py_tiler.py script packaged in PHYLUCE for each of the four to eight taxa selected for probe design (Table 1) (Faircloth *et al*. 2012; Faircloth 2015). Probes were mapped against the consensus sequences of the alignments from Hughes *et al*. (2018) in Geneious Pro v8.1 (http://www.geneious.com) to examine the distribution of probes across loci. Visual inspection of the distribution of probes initially revealed highly uneven coverage of probes across longer loci. To have the probes cover the reference alignments more evenly, we applied a staggering strategy by tiling probes every 20 bp across the locus for each of the four to eight taxa (Table 1), so that probes from the first species spanned the first 0-120 bp, and probes from the second species spanned from 20-140, and so on. This strategy ultimately improved tiling density and resulted in more even coverage for longer loci *in silico*. The probe staggering design was generated via custom scripts (Jake Enk, Arbor Biosciences). Probes that had more than 25% repeats detected on the RepeatMasker.org database were eliminated. Probes were filtered for potential self-hybridization. Each of our eight probe sets was designed with a MyBaits1 custom probe set with approximately 20,000 biotinylated probes for each set (Arbor Biosciences, Ann Arbor, Michigan).

### Mitochondrial Probes

In addition to exon probes, we designed and synthesized a separate, fish universal bait set to capture four of the most popular mitochondrial DNA (mtDNA) gene markers used in fish systematics: COI (cytochrome c oxidase subunit I), CYTB (cytochrome *b*), and 12S and 16S ribosomal DNA. The goal of maintaining separate mtDNA and nDNA probe sets is to equilibrate nDNA/mtDNA sample ratios by applying spiking dilutions of the mtDNA probe set after capturing the exon markers (mtDNA:nDNA dilution ratios = 1:1000). Probes for these four mtDNA genes were individually designed using mtDNA genomes or single sequences from NCBI that span the diversity of ray-finned fishes (*Amia calva*, *Danio rerio*, *Elops saurus*, *Harengula clupeola, Harengula jaguana*, *Oryzias latipes*, *Osteoglossum bicirrhosum*, *Polypterus ansorgii*, *Salmo trutta*, *Takifugu vermicularis*, and *Zeus faber*). A total of 7,000 oligonucleotide baits (120 bp long) tiling over the mtDNA genes with 2x density were designed using the py_tiler.py script (Faircloth *et al*. 2012; Faircloth 2015). We did not target other high-copy genes like 28S rDNA, which may have required an additional probe set and spiking dilution.

### Library Preparation and Sequencing

Eight fish species were newly sequenced for each bait set (Table 2; 56 total species sequenced). DNA extractions were performed on the GenePrep (Autogen Inc.) following manufacturer’s instructions at the Laboratory of Analytical Biology at the Smithsonian Institution National Museum of Natural History in Washington, D.C. DNA was eluted in 50 μL of Autogen R9 Buffer. Quality control was performed by running 1 μL of eluted DNA on a 1.0% agarose gel stained with GelRed (Biotium) and visually inspecting whether bands of high molecular weight DNA were visible. Library preparation was performed at Arbor Biosciences in Ann Arbor, Michigan, using a dual-round capture protocol (Li *et al*. 2013), with an 8-plex capture design. Paired-end sequencing of 100 bp reads was performed at the University of Chicago Genomics Facility on a HiSeq 4000. Samples were multiplexed with 192 in a lane, with sequencing runs containing samples for other projects not used here.

**Table 2.**
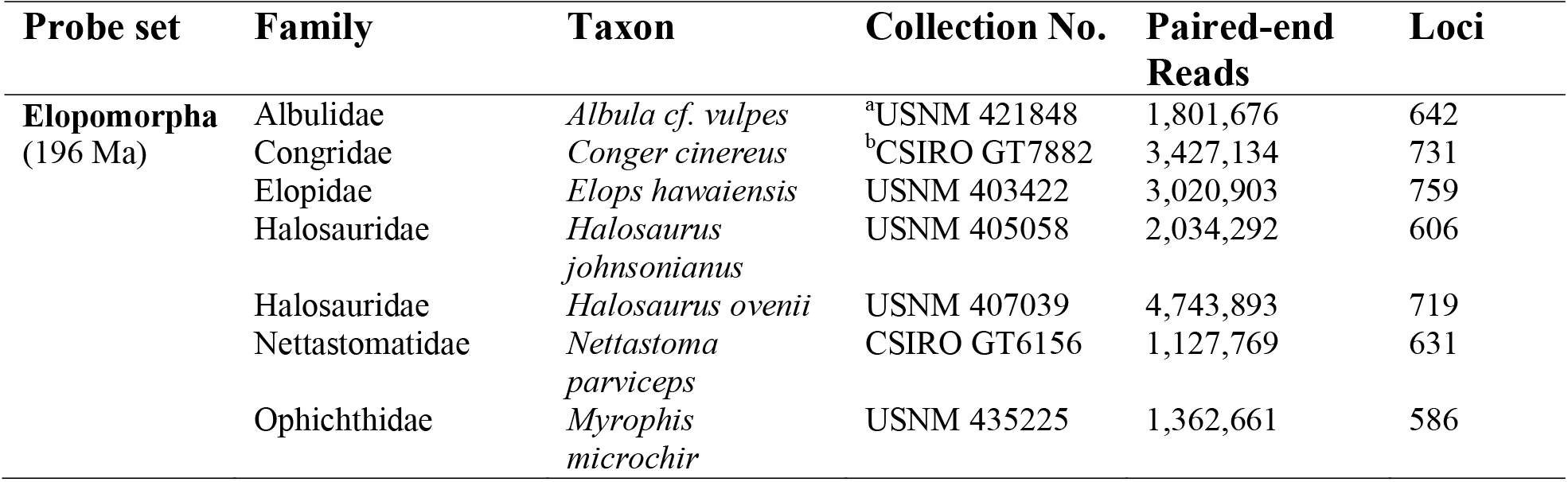
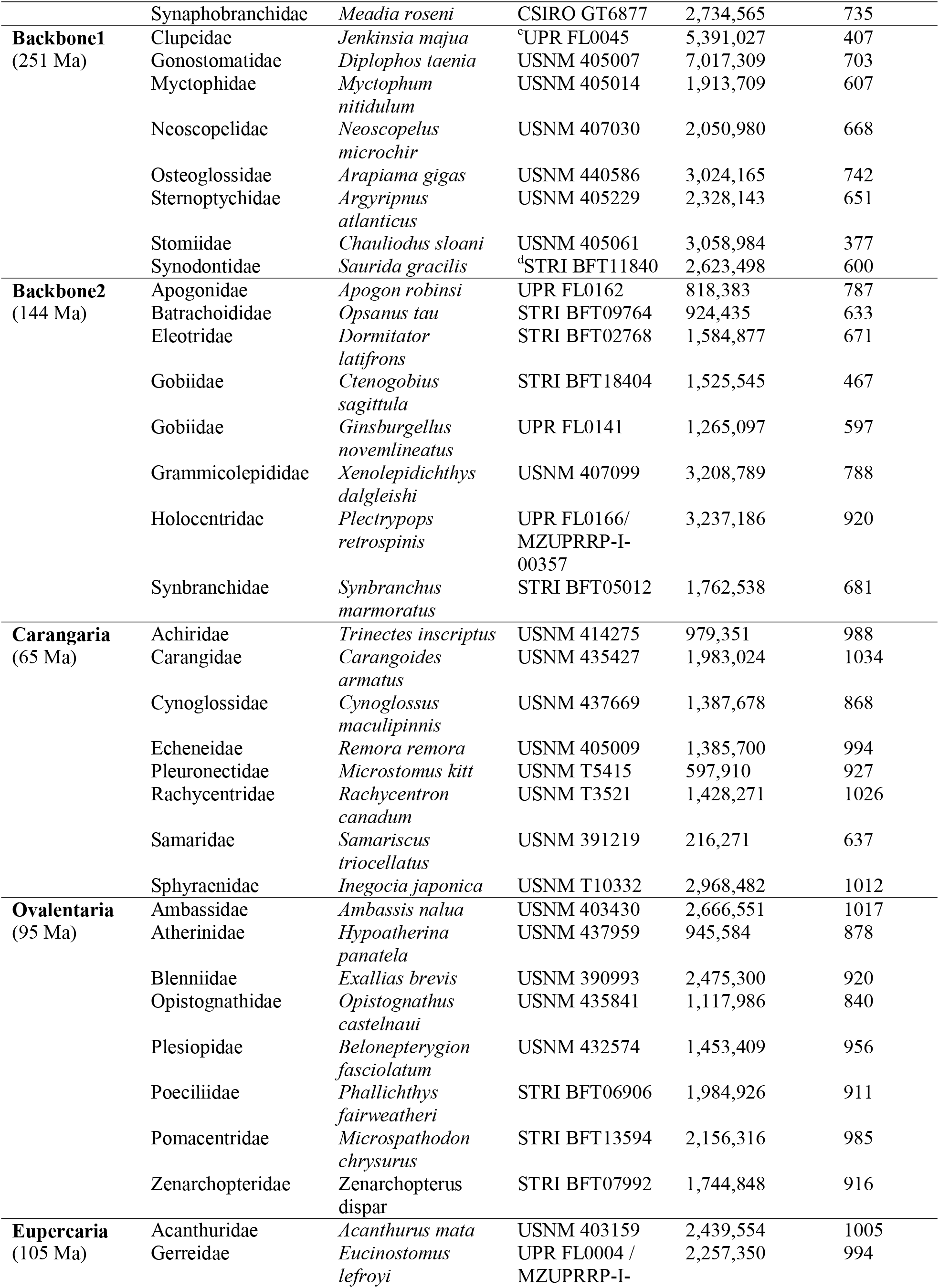
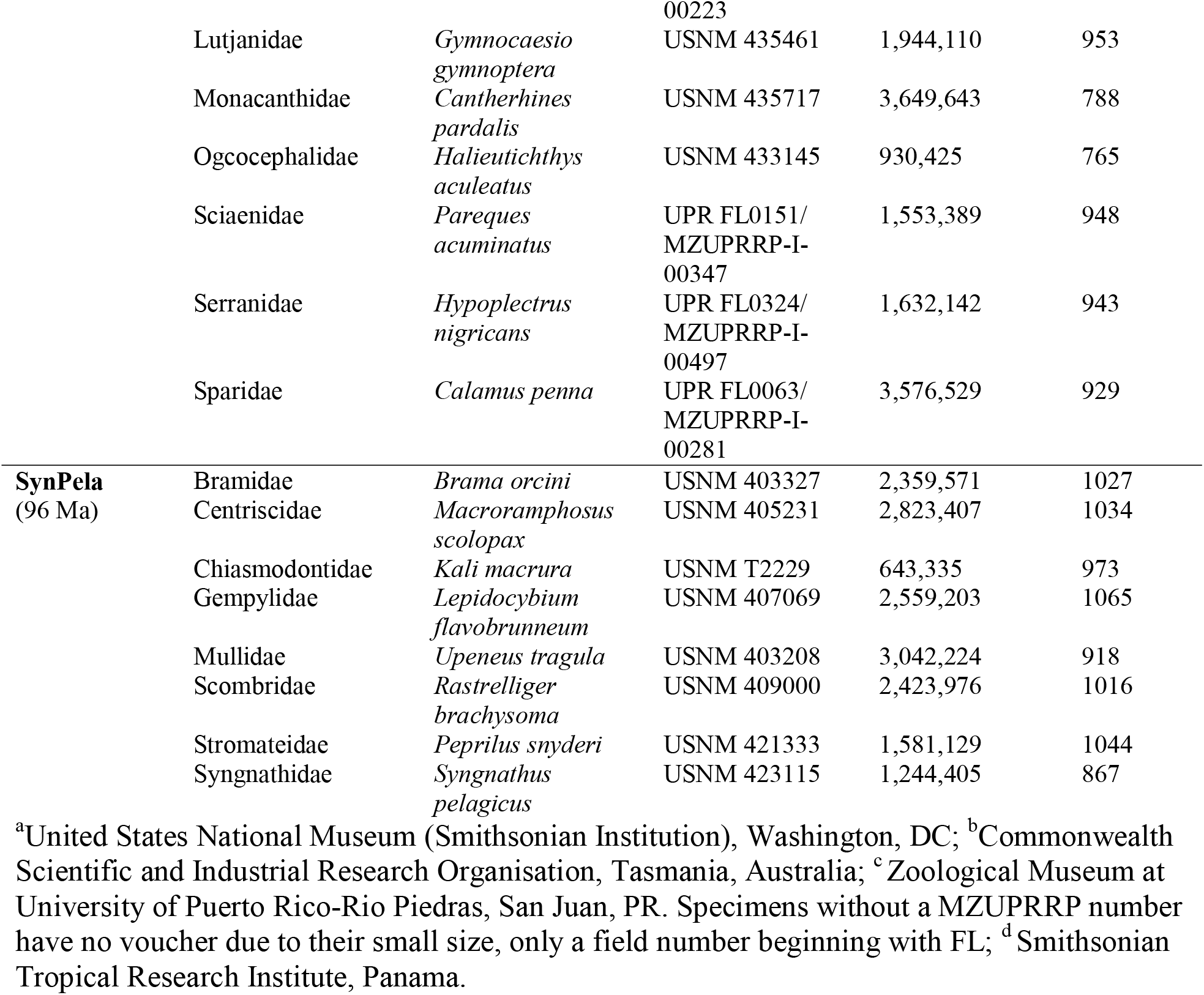
Species sequenced for each probe set and the number of reads and loci assembled. Estimated clade ages (Hughes *et al*. 2018) are indicated.

### Bioinformatics Pipeline for Exon Assembly

We developed a bioinformatics pipeline based around the software aTRAM 2.0 (Allen *et al*. 2017) with five major steps before multiple sequence alignment (Figure 4). Raw FASTQ files were quality trimmed with Trimmomatic v0.36 (Bolger *et al*. 2014), removing low quality sequences and adapter contamination with the parameters “ILLUMINACLIP:TruSeq3-PE.fa:2:30:10 LEADING:5 TRAILING:5 SLIDINGWINDOW:4:15 MINLEN:31”. Trimmed reads were then mapped against a master file containing all sequences used for bait design for any of the seven probe sets using BWA-MEM (Li & Durbin 2009). SAMtools v1.8 was used to remove optical PCR duplicates and separate the reads that mapped to each of the exons (Li *et al*. 2009b). Mapped reads were then assembled individually by exon using Velvet (Zerbino & Birney 2008), and the longest contig produced by Velvet was used as a reference sequence for aTRAM v2.0 (Allen *et al*. 2017) to extend contigs, using Trinity v2.8.5 as the assembler (Grabherr *et al*. 2011). Redundant contigs with 100% identity produced by aTRAM were removed with CD-HIT v4.8.1 using CD-HIT-EST (Li & Godzik 2006; Fu *et al*. 2012). Open reading frames for remaining contigs were identified with Exonerate (Slater & Birney 2005), using a reference sequence checked by eye for each exon, and any contigs that did not contain the open reading frame were filtered out. If only a single contig contained the open reading frame, the exon passed all filters and was used for multiple sequence alignment. If multiple contigs contained the open reading frame, the reading frames were compared with CD-HIT-EST, using a 99% identity threshold to account for potential allelic variation. If the comparison with CD-HIT-EST resulted in a single contig, that contig passed filters and was used for phylogenetic analysis; more divergent sequences were flagged and excluded from downstream analysis. Unlike another tool recently published to assemble exon-capture data, ASSEXON (Yuan *et al*. 2019), our pipeline is fully automated and does not require Geneious as part of the assembly (https://github.com/lilychughes/FishLifeExonCapture).

**Figure 4:**
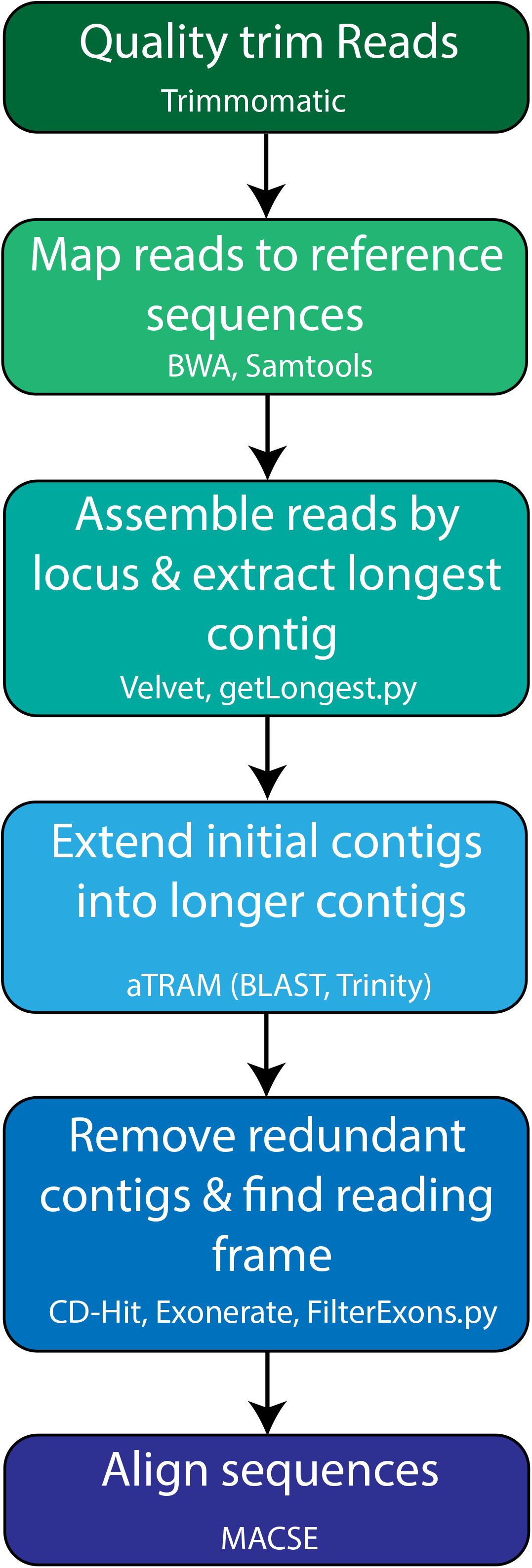
Major steps of bioinformatics pipeline and software used to transform data from Illumina reads to multiple sequence alignments. Scripts and pipeline tutorial are available on GitHub (https://github.com/lilychughes/FishLifeExonCapture).

### Alignment and Phylogenomic Analysis

Target-capture data were combined with genomic and transcriptomic data from Hughes *et al*. (2018), and 36 additional recently published genomes that were mined for orthologous exon sequences using nhmmer (Wheeler & Eddy 2013). Sequences for each exon were aligned with MACSE v2.03 (Ranwez *et al*. 2018) after cleaning out potentially non-homologous fragments with the -*cleanNonHomologousSequences* option. Alignment edges composed of more than 60% missing data as well as insertions that occurred only in a single taxon were removed with custom scripts (AlignmentCleaner.py, https://github.com/lilychughes/FishLifeExonCapture). A total of 1,104 nuclear exons filtered for paralogs were concatenated using geneStitcher.py (https://github.com/ballesterus/Utensils).

A concatenated protein matrix was analyzed under maximum likelihood (ML) with IQ-TREE v1.6.9 (Nguyen *et al*. 2015), using the best-fitting model for the entire matrix as determined using ModelFinder (Kalyaanamoorthy *et al*. 2017). A concatenated nucleotide matrix was partitioned into first, second, and third codon positions, with the best-fitting model applied to each partition. Concatenated matrices used contained only the 1,104 loci that have been filtered for paralogy; the legacy markers that failed to pass this filter in Hughes *et al*. (2018) were excluded from these analyses.

### Paralogs in Legacy Markers

Eighteen nuclear markers that have been commonly used with Sanger sequencing methods for fish phylogenetics were re-included in our probes to better connect novel sequence capture data with extensive existing datasets, but these genes had been previously filtered out for suspected paralogs. For the 18 legacy markers that were re-included (TBR1, MYH6, KIAA1239, PLAGL2, PTCHD1, RIPK4, SH3PX3, SIDKEY, SREB2, ZIC1, SVEP1, GPR61, IRBP, RNF213, Rhodopsin, SLC10A3, UBE3A, and UBE3A-like), we integrated our newly sequenced data with the matrices from Betancur-R. *et al*. (2013) for TBR1, MYH6, KIAA1239, PLAGL2, PTCHD1, RIPK4, SH3PX3, SIDKEY, SREB2, ZIC1. For the remaining genes, we pulled a selection of sequences from GenBank for SVEP1, GPR61, IRBP, RNF213, Rhodopsin, SLC10A3, UBE3A, and UBE3A-like to align with our new data. All GenBank accession numbers can be found on the sequence labels of these gene trees available on FigShare (Hughes *et al*. 2020). We inferred gene trees in IQ-TREE 1.6.9, partitioning by codon position and using the ModelFinder algorithm to determine the best-fitting sequence model. Target-capture-derived sequences falling out in unexpected positions or clades were BLASTed against the NCBI nucleotide database to determine their identity.

## Results

### Capture Efficiency of Nuclear Exons

The number of reads per sample varied substantially, from 216,271 to 7,017,309 (Table 2). However, the number of reads per sample was not correlated with the number of loci assembled per species (r^2^=0.0043; *p*=0.27). Capture efficiency (measured as the average number of exons assembled per species) varied across probe sets and across samples (Figure 5A), showing a strong negative correlation with clade (or paraphyletic “grade”) age (r^2^=0.82; *p*=0.003131; Figure 5B). Two probe sets designed to capture markers in paraphyletic groups that include more anciently diverging lineages (~251-144 Ma) with larger phylogenetic diversity (Backbone 1 and 2; Figures 1-2) tended to have lower capture efficiency (Table 2; Figure 5A), with an average of 52% of the loci captured for Backbone 1 and 61% on average captured for Backbone 2. This was also the case for the rather ancient Elopomorpha clade (196 Ma), for which samples sequenced had only a 60% capture rate. Probe sets designed for more recently diverged percomorph clades (~105-66 Ma) had much higher numbers of loci assembled on average, with 83% for Carangaria, 82% for Ovalentaria, 81% for Eupercaria, and 89% for the Syngnatharia + Pelagiaria clade. Probe sets are publicly available at Arbor Biosciences, Ann Arbor, MI. Exon alignments ranged from a minimum of 60% up to 93.9% sequence identity across the eight model genomes originally used to discover single-copy markers, but the legacy markers tended to have >80% sequence identity (Figure 5C).

**Figure 5:**
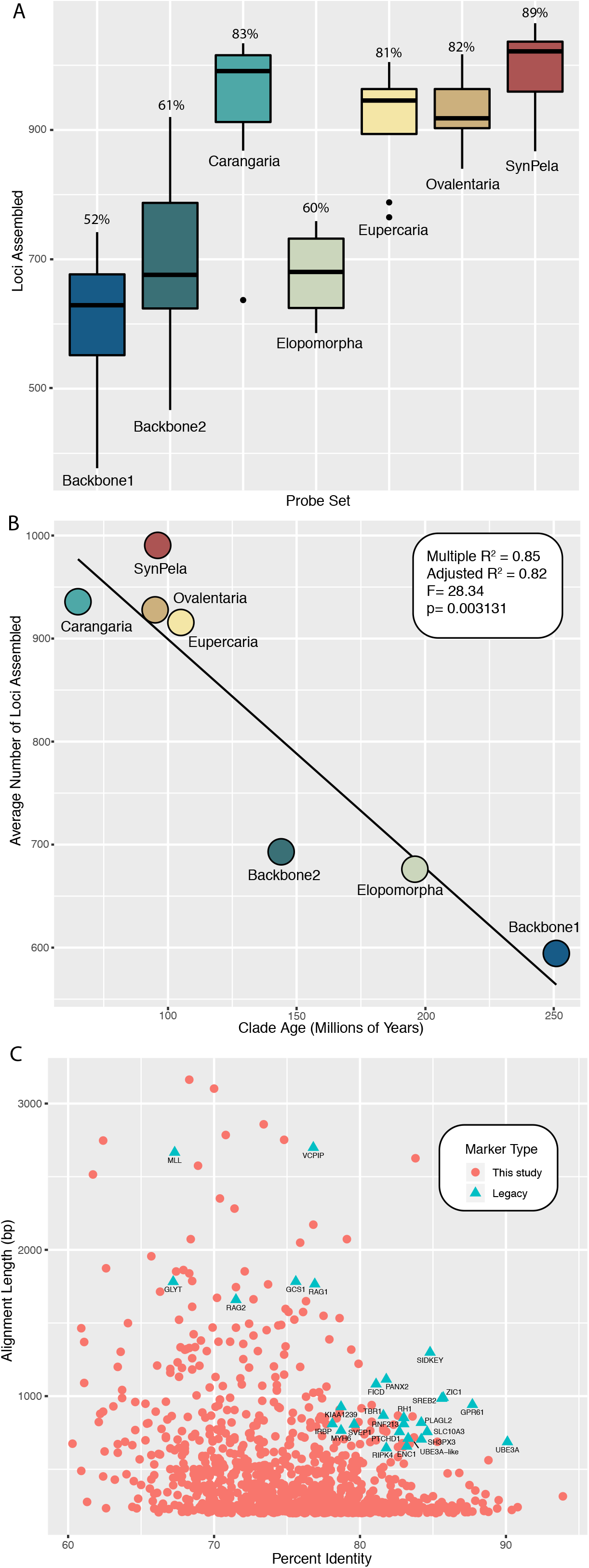
A) Box-and-whisker plots showing the variation in the number of exons captured for each probe set. Eight newly sequenced taxa were used for each probe set. Percentages above each box refer to the average percentage of exons assembled by probe set. B) The average number of loci captured for each of the seven probe sets plotted against the age of the most recent common ancestor of that particular group, showing higher capture of loci for younger clades. C) Percent sequence identity and length across alignments of eight model genomes (*L. oculatus, A. japonica, D. rerio, G. morhua, O. latipes, O. niloticus, G. aculeatus*, and *T. nigroviridis*) for exons presented in this study (pink circles) compared to legacy markers (blue triangles).

### Mitochondrial Gene Capture

Mitochondrial genes for which we designed probes (12S, 16S, COI, ATPase6, and CYTB) tended to have the best rate of capture. Complete sequences of 12S, 16S, and COI assembled for all taxa, while ND6, for which we did not design probes, was only represented for 29 taxa, the lowest of any mitochondrial coding gene. CYTB, which was included in the probe set, assembled for 56 of 58 total taxa, but ATPase6 had a relatively poor capture rate, only assembling for 36 species.

### Phylogenomic Analysis

Combining the new data for 56 samples collected through exon-capture plus 38 additional recently published fish genomes with the dataset of Hughes *et al*. (2018) generated a matrix with 394 taxa representing all major groups of ray-finned fishes with three lobe-finned (sarcopterygian) outgroups, with a final length of 549,861 bp (183,287 amino acid sites). The entire matrix had 72% present data, excluding loci suspected of having paralogs (Table 3). The average locus alignment length for genes included in the matrix was 499 bp (range: 129-5,055 bp), and the average number of parsimony-informative sites per locus was 340 (range: 75-3,435).

**Table 3.** Legacy markers and their paralogs assembled from exon-capture data.

| Locus ID | Gene Name | Paralogs Assembled |
| --- | --- | --- |
| E1541 | TBR1 | - |
| E1728 | KIAA1239 | - |
| E1730 | MYH6 | MYH7, MYH6-like, MYHb |
| E1732 | ENC1 | ENC2 |
| E1735 | PLAGL2 | PLAGL1 |
| E1736 | PTCHD1 | (Failed to assemble) |
| E1737 | RIPK4 | - |
| E1738 | SH3PX3 (SNX33) | SNX18 |
| E1739 | SIDKEY | - |
| E1740 | SREB2 (GPR85) | GPR173 |
| E1741 | ZIC1 | ZIC4 |
| E1746 | SVEP1 | - |
| E1747 | GPR61 | - |
| E1748 | IRBP | - |
| E1749 | RNF213 | RNF213b-like, RNF213a-like |
| E1750 | RH | EXORH |
| E1751 | SLC10A3 | - |
| E1752 | UBEA3 | - |
| E1753 | UBEA3-like | UBEA3 |

ModelFinder selected JTT+I+F+G for the protein matrix, and GTR+I+F+G for the first two codon positions, with TVM+I+F+G for the third codon position. Both topologies inferred with IQ-TREE matched previous results obtained by Hughes *et al*. (2018), with newly added taxa placed in their expected phylogenetic placements (Figures 1-3). Tree files and phylogenomic matrices are available on FigShare (Hughes *et al*. 2020).

### Paralogs in Legacy Markers

We examined gene trees by eye for evidence paralogs had been assembled for 19 markers that were included in our probe set, despite failing previous tests to exclude loci with paralogs. Nine of these loci assembled one or more paralogs when we applied our new pipeline on the raw reads obtained with sequence capture instead of a single orthologous locus (Table 3). One locus (PTCHD1) could only be assembled for seven of the 56 newly sequenced samples, which made the paralogy assessment difficult. Annotation of the paralogous sequences was determined by blasting assembled contigs against the NCBI nucleotide database.

## Discussion

### Deep Probe Sets Capture Across Deep Phylogenetic Divergences

We present resources for capturing conserved exon sequences for all groups of teleost fishes, including probe sets for early branching lineages (“Backbone 1”), Elopomorpha, Acanthomorphata (“Backbone 2”), and multiple major percomorph radiations including Syngnatharia + Pelagiaria, Carangaria, Ovalentaria, and Eupercaria. The exon markers presented here have been explicitly tested for orthology using a large database of 303 bony fish species (Hughes *et al*. 2018), and have been screened for paralogs derived from ancient vertebrate or teleost-specific whole genome duplication events. Capture efficiency is strongly correlated with the phylogenetic span of taxa used to design the probes, with probe sets designed more specifically for younger (~105-66 Ma) percomorph clades (Syngnatharia-Pelagiaria, Carangaria, Ovalentaria, Eupercaria) capturing 200-300 more loci on average than those designed for more ancient (~251-144 Ma) and/or taxonomically disparate clades (Elopomorpha and Backbones 1 and 2; Figure 5B). Estimates for the divergence times of major percomorph series vary, but the youngest estimates place their origin near the Cretaceous-Paleogene boundary, 66 million years ago (Alfaro *et al*. 2018). Conversely, the paraphyletic taxonomic groups spanned by Backbone 1 diverged in the Permian or Triassic, and the late Jurassic to early Cretaceous for Backbone 2 (Betancur-R. *et al*. 2013; Near *et al*. 2013a; Hughes *et al*. 2018). The larger number of nucleotide substitutions accumulated across older clades causes the probes to have less affinity for the targeted DNA regions *in vitro*, and we noticed a substantial increase in the number of loci captured for those clades younger than 100 million years (Figure 5B). One way to improve capture efficiency for particular projects would be to select more closely related taxa to design a new (more specific) probe set.

Despite the variation in the number of loci assembled, all samples with exon-capture data were resolved in their expected clades, and the ML topologies at major nodes matched that of the ML topologies of Hughes *et al*. (2018). One family, Clupeidae, was not monophyletic, but this result has been reported before in previous phylogenetic analyses (Betancur-R *et al*. 2017), and may reflect the need of taxonomic revision or poor taxonomic sampling rather than underlying phylogenetic estimation error arising from the exon-capture data. These markers appear to be informative for deep divergences in fishes, and the backbone of the ray-finned fish tree largely matches inferences based on legacy gene markers (Near *et al*. 2012; Betancur-R. *et al*. 2013), though many areas of the tree have only been investigated with sparse taxonomic sampling and will require more thorough investigation with additional sequencing. While deep divergences are the focus of this paper, conserved exon markers have also been shown to contain information appropriate for shallower divergences at the phylogeographic level (Rincon-Sandoval *et al*. 2019). The flanking intron regions, which are highly variable, have been removed for the analyses presented here, but we include a branch of our bioinformatic pipeline to additionally use the flanking intron sequences for projects with a more recent evolutionary focus.

### Exon markers can be integrated with existing and future datasets

Taxonomic sampling is critical for accurate phylogenetic analysis (Heath *et al*. 2008; Betancur-R. *et al*. 2019), and sequence capture methods are a cost-effective approach for increasing taxonomic sampling across a large number of loci (Lemmon & Lemmon 2013). But both whole-genome sequences (Malmstrøm *et al*. 2016; Musilova *et al*. 2019) and transcriptome sequences (e.g. Hughes *et al*. 2018; Dai *et al*. 2018) are becoming available for an increasing diversity of fish species. These exon markers can be easily mined from public transcriptome or genome data as they become available, increasing taxonomic sampling for the group of interest without duplicating sequencing effort. Taxonomically dense super-matrix approaches in fishes (e.g. Rabosky *et al*. 2018) primarily rely on exon sequences deposited in NCBI. Currently, there are more than 20,000 sequences of RAG1 for teleosts available in NCBI (as of March 20, 2019), more than 35,000 teleost rhodopsin sequences, and even larger numbers for mitochondrial genes like CYTB (>130,000 sequences). This is a rich resource that can be combined with exon capture data for the probe sets described here to reduce missing data that are often rampant in super matrix approaches but still produce taxonomically dense trees.

### Paralogs in sequence capture datasets

Many nuclear exon fragments that have been in wide use in fish phylogenetics for more than a decade do not appear to have paralogs, and new sequence capture data could be easily integrated from genes like RAG1, RAG2, PANX2, MLL, VCPIP, GLYT, GCS1, and FICD. Many of these exons were defined as ‘single-copy’ based on the comparisons of the relatively few fish genomes available at the time (Li *et al*. 2007). But the specificity of primers designed for nested PCR approaches to amplify and sequence these loci has been a successful strategy to obtain orthologous genes for phylogenetic inference in fishes (Li *et al*. 2007, 2008, 2010; Near *et al*. 2012, 2013b; Wainwright *et al*. 2012; Betancur-R. *et al*. 2013). Shotgun sequencing of enriched libraries, in contrast, is a more challenging approach for assembling orthologous genes, since sequence reads of two or more paralogous copies may be sequenced by this approach and need to be separated using bioinformatic pipelines. Nineteen legacy markers included in our probe set previously had been excluded from downstream phylogenetic analyses due to paralogy issues detected either by comparing additional genomes or by topology tests of gene trees (Table 3). Due to high similarity in certain parts of the coding region to the reference coding sequence used in Exonerate, more than half of these assembled paralogous loci passed through to the alignment stage. Often it was only the paralog that was assembled, and the assembly of multiple contigs was not a reliable way to flag paralogy. The pipeline implemented here (Figure 4) attempts to remove redundant contigs with CD-HIT at a 99% similarity threshold across the reading frame when multiple contigs assemble, but exons that fail this test are not passed onto the alignment stage. Paralogs of ENC1, MYH6, ZIC1 and other genes known to be duplicated (Table 3) passed on to the alignment stage, and do not appear to have assembled the orthologous sequence.

However, a majority of the sequences assembled orthologous exons. With additional scrutiny for paralogs using gene trees, these data are still quite useful for integration with older datasets.

## Acknowledgements

This research was supported by National Science Foundation (NSF) grants NSF-DEB-1929248 and NSF-DEB-1932759 to R.B.R., NSF-DEB-1541554 and NSF-DEB-1457426 to G.O., and NSF-DEB-1541552 to C.C.B. We are grateful to Jake Enk (Arbor Biosciences) for his assistance with probe design. We thank Rose Peterson and Victoria Rodriguez for assistance with laboratory work, and Diane Pitassy for access to tissues at the USNM. All data processing and phylogenetic analysis were conducted on the Pegasus HPC cluster at George Washington University.

## Data Accessibility

Raw reads for newly sequenced exon-capture data are archived on NCBI under Bioproject number PRJNA605876. Newick files and phylogenomic matrices are available on FigShare (doi: 10.6084/m9.figshare.11844783), and pipeline scripts to analyze data and a tutorial are available on GitHub (https://github.com/lilychughes/FishLifeExonCapture). The protein tree topology will be made available on Open Tree of Life.

## Author Contributions

L.C.H., R.B-R., G.O., K.A.C, C.L., and D.A. contributed to the design of the study. R.B-R., C.C.B., and W.W. provided tissues. L.C.H., H.S., C.L., and R.B-R. analyzed the data. L.C.H., G.O., and R.B-R. wrote the paper with input from all authors.

## Notes

doi:10.6084/m9.figshare.11844783

## References

Abascal F, Zardoya R, Telford MJ (2010) TranslatorX: multiple alignment of nucleotide sequences guided by amino acid translations. Nucleic Acids Research, 38, W7–W13.

Alfaro ME, Faircloth BC, Harrington RC et al. (2018) Explosive diversification of marine fishes at the Cretaceous-Palaeogene boundary. Nature Ecology & Evolution.

Allen JM, Boyd B, Nguyen N et al. (2017) Phylogenomics from Whole Genome Sequences Using aTRAM. Systematic Biology, 66, 786–798.

Arcila D, Ortí G, Vari R et al. (2017) Genome-wide interrogation advances resolution of recalcitrant groups in the tree of life. Nature Ecology & Evolution, 1, 0020.

Betancur-R. R, Broughton RE, Wiley EO et al. (2013) The Tree of Life and a New Classification of Bony Fishes. PLoS Currents, 5.

Betancur-R R, Orti G, O Wiley E et al. (2017) Phylogenetic classification of bony fishes. BMC Evolutionary Biology, 1–40.

Betancur-R. R, Arcila D, Vari RP et al. (2019) Phylogenomic incongruence, hypothesis testing, and taxonomic sampling: The monophyly of characiform fishes. Evolution, 73, 329–345.

Bolger AM, Lohse M, Usadel B (2014) Trimmomatic: a flexible trimmer for Illumina sequence data. Bioinformatics, 30, 2114–2120.

Braasch I, Peterson SM, Desvignes T et al. (2015) A new model army: Emerging fish models to study the genomics of vertebrate Evo-Devo. Journal of Experimental Zoology Part B: Molecular and Developmental Evolution, 324, 316–341.

Bybee SM, Bracken-Grissom H, Haynes BD et al. (2011) Targeted Amplicon Sequencing (TAS): A Scalable Next-Gen Approach to Multilocus, Multitaxa Phylogenetics. Genome Biology and Evolution, 3, 1312–1323.

Chakrabarty P, Faircloth BC, Alda F et al. (2017) Phylogenomic Systematics of Ostariophysan Fishes: Ultraconserved Elements Support the Surprising Non-Monophyly of Characiformes. Systematic Biology, 66, 881–895.

Chen W-J, Bonillo C, Lecointre G (2003) Repeatability of clades as a criterion of reliability: a case study for molecular phylogeny of Acanthomorpha (Teleostei) with larger number of taxa. Molecular Phylogenetics and Evolution, 26, 262–288.

Dai W, Zou M, Yang L et al. (2018) Phylogenomic Perspective on the Relationships and Evolutionary History of the Major Otocephalan Lineages. Scientific Reports, 8, 205.

Dehal P, Boore JL (2005) Two Rounds of Whole Genome Duplication in the Ancestral Vertebrate (P Holland, Ed,). PLoS Biology, 3, e314.

Dettai A, Lecointre G (2005) Further support for the clades obtained by multiple molecular phylogenies in the acanthomorph bush. Comptes Rendus - Biologies, 328, 674–689.

Dettaï A, Lecointre G (2008) New insights into the organization and evolution of vertebrate IRBP genes and utility of IRBP gene sequences for the phylogenetic study of the Acanthomorpha (Actinopterygii: Teleostei). Molecular Phylogenetics and Evolution, 48, 258–269.

Dornburg A, Townsend JP, Brooks W et al. (2017) New Insights on the Sister Lineage of Percomorph Fishes with an Anchored Hybrid Enrichment Dataset. Molecular Phylogenetics and Evolution, 110, 27–38.

Edwards S V., Cloutier A, Baker AJ (2017) Conserved Nonexonic Elements: A Novel Class of Marker for Phylogenomics. Systematic Biology, 66, 1028–1044.

Eytan RI, Evans BR, Dornburg A et al. (2015) Are 100 enough? Inferring acanthomorph teleost phylogeny using Anchored Hybrid Enrichment. BMC evolutionary biology, 15, 113.

Faircloth BC (2015) PHYLUCE is a software package for the analysis of conserved genomic loci. Bioinformatics, 32, 786–788.

Faircloth BC, Alda F, Hoekzema K et al. (2020) A target enrichment bait set for studying relationships among ostariophysan fishes. Copeia, 108, 47–60.

Faircloth BC, McCormack JE, Crawford NG et al. (2012) Ultraconserved elements anchor thousands of genetic markers spanning multiple evolutionary timescales. Systematic Biology, 61, 717–726.

Faircloth BC, Sorenson L, Santini F, Alfaro ME (2013) A Phylogenomic Perspective on the Radiation of Ray-Finned Fishes Based upon Targeted Sequencing of Ultraconserved Elements (UCEs). PLoS One, 8, e65923.

Friedman M, Feilich KL, Beckett HT et al. (2019) A phylogenomic framework for pelagiarian fishes (Acanthomorpha: Percomorpha) highlights mosaic radiation in the open ocean. Proceedings of the Royal Society B: Biological Sciences, 286, 20191502.

Fu L, Niu B, Zhu Z, Wu S, Li W (2012) CD-HIT: accelerated for clustering the next-generation sequencing data. Bioinformatics, 28, 3150–3152.

Grabherr MG, Haas BJ, Yassour M et al. (2011) Full-length transcriptome assembly from RNA-Seq data without a reference genome. Nature biotechnology, 29, 644–652.

Harrington RC, Faircloth BC, Eytan RI et al. (2016) Phylogenomic analysis of carangimorph fishes reveals flatfish asymmetry arose in a blink of the evolutionary eye. BMC Evolutionary Biology, 16, 224.

Heath TA, Hedtke SM, Hillis DM (2008) Taxon sampling and the accuracy of phylogenetic analyses. Journal of Systematics And Evolution, 46, 239–257.

Hughes LC, Ortí G, Huang Y et al. (2018) Comprehensive phylogeny of ray-finned fishes (Actinopterygii) based on transcriptomic and genomic data. Proceedings of the National Academy of Sciences, 115, 6249–6254.

Hughes LC, Ortí G, Saad H et al. (2020) Data from: Exon probe sets and bioinformatics pipelines for all levels of fish phylogenomics. FigShare, doi:10.6084/m9.figshare.11844783.

Ilves KL, López-Fernández H (2014) A targeted next-generation sequencing toolkit for exon-based cichlid phylogenomics. Molecular Ecology Resources, 14, 802–811.

Ilves KL, Torti D, López-Fernández H (2017) Exon-based phylogenomics strengthens the phylogeny of Neotropical cichlids and identifies remaining conflicting clades (Cichliformes: Cichlidae: Cichlinae). Molecular Phylogenetics and Evolution, 118, 232–243.

Inoue J, Sato Y, Sinclair R, Tsukamoto K, Nishida M (2015) Rapid genome reshaping by multiple-gene loss after whole-genome duplication in teleost fish suggested by mathematical modeling. Proceedings of the National Academy of Sciences, 112, 14918–14923.

Irisarri I, Singh P, Koblmüller S et al. (2018) Phylogenomics uncovers early hybridization and adaptive loci shaping the radiation of Lake Tanganyika cichlid fishes. Nature Communications, 9, 3159.

Jiang J, Yuan H, Zheng X et al. (2019) Gene markers for exon capture and phylogenomics in ray finned fishes. Ecology and Evolution, 9, 3973–3983.

Kalyaanamoorthy S, Minh BQ, Wong TKF, von Haeseler A, Jermiin LS (2017) ModelFinder: fast model selection for accurate phylogenetic estimates. Nature Methods, 14, 587–589.

Kuang T, Tornabene L, Li J et al. (2018) Phylogenomic analysis on the exceptionally diverse fish clade Gobioidei (Actinopterygii: Gobiiformes) and data-filtering based on molecular clocklikeness. Molecular Phylogenetics and Evolution, 128, 192–202.

Lemmon AR, Emme SA, Lemmon EM (2012) Anchored Hybrid Enrichment for Massively High-Throughput Phylogenomics. Systematic Biology, 61, 727–744.

Lemmon EM, Lemmon AR (2013) High-Throughput Genomic Data in Systematics and Phylogenetics. Annual Review of Ecology, Evolution, and Systematics, 44, 99–121.

Li C, Corrigan S, Yang L et al. (2015) DNA capture reveals transoceanic gene flow in endangered river sharks. Proceedings of the National Academy of Sciences, 112, 13302–13307.

Li B, Dettaï A, Cruaud C et al. (2009a) RNF213, a new nuclear marker for acanthomorph phylogeny. Molecular Phylogenetics and Evolution, 50, 345–363.

Li H, Durbin R (2009) Fast and accurate short read alignment with Burrows-Wheeler transform. Bioinformatics, 25, 1754–1760.

Li W, Godzik A (2006) Cd-hit: a fast program for clustering and comparing large sets of protein or nucleotide sequences. Bioinformatics, 22, 1658–1659.

Li H, Handsaker B, Wysoker A et al. (2009b) The Sequence Alignment/Map format and SAMtools. Bioinformatics, 25, 2078–2079.

Li C, Hofreiter M, Straube N, Corrigan S, Naylor GJP (2013) Capturing protein-coding genes across highly divergent species. BioTechniques, 54, 321–326.

Li C, Lu G, Ortí G (2008) Optimal Data Partitioning and a Test Case for Ray-Finned Fishes (Actinopterygii) Based on Ten Nuclear Loci (T Buckley, Ed,). Systematic Biology, 57, 519–539.

Li C, Ortí G, Zhang G, Lu G (2007) A practical approach to phylogenomics: the phylogeny of ray-finned fish (Actinopterygii) as a case study. BMC Evolutionary Biology, 7, 44.

Li C, Ortí G, Zhao J (2010) The phylogenetic placement of sinipercid fishes (“ Perciformes”) revealed by 11 nuclear loci. Molecular Phylogenetics and Evolution, 56, 1096–1104.

Li C, Ricardo B-R, Leo Smith W, Ortí G (2011) Monophyly and interrelationships of Snook and Barramundi (Centropomidae sensu Greenwood) and five new markers for fish phylogenetics. Molecular Phylogenetics and Evolution, 60, 463–471.

Li C, Riethoven JJM, Naylor GJP (2012) EvolMarkers: A database for mining exon and intron markers for evolution, ecology and conservation studies. Molecular Ecology Resources, 12, 967–971.

Longo SJ, Faircloth BC, Meyer A et al. (2017) Phylogenomic analysis of a rapid radiation of misfit fishes (Syngnathiformes) using ultraconserved elements. Molecular Phylogenetics and Evolution, 113, 33–48.

Lopez JA, Chen W-J, Ortí G (2004) Esociform Phylogeny. Copeia, 3, 449–464.

Lovejoy NR, Iranpour M, Collette BB (2004) Phylogeny and jaw ontogeny of beloniform fishes. Integrative and Comparative Biology, 44, 366–377.

Malmstrøm M, Matschiner M, Tørresen OK et al. (2016) Evolution of the immune system influences speciation rates in teleost fishes. Nature Genetics, 48, 1204–1210.

Musilova Z, Cortesi F, Matschiner M et al. (2019) Vision using multiple distinct rod opsins in deep-sea fishes. Science, 364, 588–592.

Near TJ, Dornburg A, Eytan RI et al. (2013a) Phylogeny and tempo of diversification in the superradiation of spiny-rayed fishes. Proceedings of the National Academy of Sciences, 110, 12738–12743.

Near TJ, Dornburg A, Eytan RI et al. (2013b) Phylogeny and tempo of diversification in the superradiation of spiny-rayed fishes. Proceedings of the National Academy of Sciences, 110, 12738–12743.

Near TJ, Eytan RI, Dornburg A et al. (2012) Resolution of ray-finned fish phylogeny and timing of diversification. Proceedings of the National Academy of Sciences, 109, 13698–13703.

Nguyen LT, Schmidt HA, Von Haeseler A, Minh BQ (2015) IQ-TREE: A fast and effective stochastic algorithm for estimating maximum-likelihood phylogenies. Molecular Biology and Evolution, 32, 268–274.

Rabosky DL, Chang J, Title PO et al. (2018) An inverse latitudinal gradient in speciation rate for marine fishes. Nature, 559, 392–395.

Ranwez V, Douzery EJP, Cambon C, Chantret N, Delsuc F (2018) MACSE v2: Toolkit for the Alignment of Coding Sequences Accounting for Frameshifts and Stop Codons. Molecular biology and evolution, 35, 2582–2584.

Ranwez V, Harispe S, Delsuc F, Douzery EJP (2011) MACSE: Multiple alignment of coding SEquences accounting for frameshifts and stop codons. PLoS ONE, 6.

Rincon-Sandoval M, Betancur-R R, Maldonado-Ocampo JA (2019) Comparative phylogeography of trans-Andean freshwater fishes based on genome-wide nuclear and mitochondrial markers. Molecular Ecology, 28, 1096–1115.

Roxo FF, Ochoa LE, Sabaj MH et al. (2019) Phylogenomic reappraisal of the Neotropical catfish family Loricariidae (Teleostei: Siluriformes) using ultraconserved elements. Molecular Phylogenetics and Evolution, 135, 148–165.

Slater GSC, Birney E (2005) Automated generation of heuristics for biological sequence comparison. BMC Bioinformatics, 6, 1–11.

Song S, Zhao J, Li C (2017) Species delimitation and phylogenetic reconstruction of the sinipercids (Perciformes: Sinipercidae) based on target enrichment of thousands of nuclear coding sequences. Molecular Phylogenetics and Evolution, 111, 44–55.

Stout CC, Tan M, Lemmon AR, Lemmon EM, Armbruster JW (2016) Resolving Cypriniformes relationships using an anchored enrichment approach. BMC Evolutionary Biology, 16, 244.

Straube N, Li C, Mertzen M, Yuan H, Moritz T (2018) A phylogenomic approach to reconstruct interrelationships of main clupeocephalan lineages with a critical discussion of morphological apomorphies. BMC Evolutionary Biology, 18, 158.

Sun Y, Huang Y, Li X et al. (2016) Fish-T1K (Transcriptomes of 1,000 Fishes) Project: large-scale transcriptome data for fish evolution studies. GigaScience, 5, 18.

Vandepoele K, De Vos W, Taylor JS, Meyer A, Van de Peer Y (2004) Major events in the genome evolution of vertebrates: Paranome age and size differ considerably between ray-finned fishes and land vertebrates. Proceedings of the National Academy of Sciences, 101, 1638–1643.

Wainwright PC, Smith WL, Price SA et al. (2012) The Evolution of Pharyngognathy: A Phylogenetic and Functional Appraisal of the Pharyngeal Jaw Key Innovation in Labroid Fishes and Beyond. Systematic Biology, 61, 1001–1027.

Wheeler TJ, Eddy SR (2013) nhmmer: DNA homology search with profile HMMs. Bioinformatics, 29, 2487–2489.

Yin G, Pan Y, Sarker A et al. (2019) Molecular systematics of Pampus (Perciformes: Stromateidae) based on thousands of nuclear loci using target-gene enrichment. Molecular Phylogenetics and Evolution, 140, 106595.

Yuan H, Atta C, Tornabene L, Li C (2019) Assexon: Assembling Exon Using Gene Capture Data. Evolutionary Bioinformatics, 15, 117693431987479.

Zerbino DR, Birney E (2008) Velvet: Algorithms for de novo short read assembly using de Bruijn graphs. Genome Research, 18, 821–829.

